# Developmental demands contribute to early neuromuscular degeneration in CMT2D mice

**DOI:** 10.1101/2020.05.21.106252

**Authors:** James N. Sleigh, Aleksandra M. Mech, Giampietro Schiavo

**Affiliations:** Department of Neuromuscular Diseases, Institute of Neurology, University College London, London WC1N 3BG, UK; UK Dementia Research Institute, University College London, London WC1E 6BT, UK; Discoveries Centre for Regenerative and Precision Medicine, University College London Campus, London WC1N 3BG, UK

**Keywords:** aminoacyl-tRNA synthetase (ARS), Charcot-Marie-Tooth disease (CMT), denervation, *GARS1*, glycyl-tRNA synthetase (GlyRS), motor neuron, neuromuscular junction (NMJ), peripheral neuropathy, synapse elimination

## Abstract

Dominantly inherited, missense mutations in the widely expressed housekeeping gene, *GARS1*, cause Charcot-Marie-Tooth type 2D (CMT2D), a peripheral neuropathy characterised by muscle weakness and wasting in limb extremities. Mice modelling CMT2D display early and selective neuromuscular junction (NMJ) pathology, epitomised by disturbed maturation and neurotransmission, leading to denervation. Indeed, the NMJ disruption has been reported in several different muscles; however, a systematic comparison of neuromuscular synapses from distinct body locations has yet to be performed. We therefore analysed NMJ development and degeneration across five different wholemount muscles to identify key synaptic features contributing to the distinct pattern of neurodegeneration in CMT2D mice. Denervation was found to occur along a distal-to-proximal gradient, providing a cellular explanation for the greater weakness observed in mutant *Gars* hindlimbs compared to forelimbs. Nonetheless, muscles from similar locations and innervated by axons of equivalent length showed significant differences in neuropathology, suggestive of additional factors impacting on site-specific neuromuscular degeneration. Defective NMJ development preceded and associated with degeneration, but was not linked to a delay of wild-type NMJ maturation processes. Correlation analyses indicate that muscle fibre type nor synaptic architecture explain the differential denervation of CMT2D NMJs, rather it is the extent of post-natal synaptic growth that predisposes to neurodegeneration. Together, this work improves our understanding of the mechanisms driving synaptic vulnerability in CMT2D and hints at pertinent pathogenic pathways.

## Introduction

Charcot-Marie-Tooth disease (CMT) is an inherited peripheral neuropathy typified by degeneration of motor and sensory neurons, which triggers progressive muscle wasting and sensory deficits mainly in the feet and hands^1^. When genetic neuropathy results from demyelination and presents with slowed nerve conduction speeds, it is classically categorised as CMT1, whereas CMT2 results from axon degeneration unrelated to myelin disruption. Mutations in more than 100 genes cause CMT, the majority of which lead to greater dysfunction in lower limbs^2^. There is thus a length-dependency to the condition, such that peripheral nerves with the longest axons are generally more affected; this suggests that cellular processes most impacted by the extreme morphology of neurons, for example axonal transport, may play an important role in CMT pathogenesis^3–5^.

Caused by dominantly inherited, missense mutations in the widely and constitutively expressed housekeeping gene, *GARS1* (Ref. 6), the 2D subtype of CMT (CMT2D) usually manifests during adolescence and, unlike most forms of the disease, frequently displays upper limb predominance^7,8^. The *GARS1*-encoded protein, glycyl-tRNA synthetase (GlyRS), charges glycine to its cognate tRNA for protein translation; yet, it is a toxic gain-of-function that likely drives neurodegeneration^9–11^. The mechanisms underlying neuronal selectivity in CMT2D remain unresolved; however, aberrant protein-protein interactions caused by relaxation of the GlyRS structure appear to underlie the neomorphic function^12,13^. Indeed, neuropathy-causing mutant GlyRS has been shown to mis-interact with the extracellular domains of neuronal transmembrane receptors, neuropilin 1 and tropomyosin receptor kinases (Trks) A-C^13,14^. Permitting this, GlyRS is secreted from several different cell types and circulates in mammalian serum, likely for a non-canonical and still only partially understood function^13,15,16^.

The neuromuscular junction (NMJ) is the specialised synapse connecting lower motor neurons to muscle fibres, and is dysfunctional in several CMT subtype models^17–22^. Indeed, CMT2D mice display loss of NMJ integrity in multiple hindlimb muscles without spinal cord motor neuron degeneration^9,23–27^. By directly comparing a proximal and a distal muscle, we have previously shown that the neuromuscular synapse is an important site of selective and early pathology^25^, replicating the muscle weakness pattern of patients. Also contributing to reduced strength and independently from denervation, these neuropathic mice display a pre-synaptic disruption in neurotransmission that correlates with disease severity and worsens with age^26^. Neuromuscular degeneration is replicated in a *Drosophila melanogaster* model for *GARS1* neuropathy and is dependent on the toxic accumulation of mutant GlyRS at the NMJ^16^. Suggestive of a non-cell autonomous mechanism, this pathological build-up requires muscle-secreted, but not neuron-derived, GlyRS, and appears to be mediated by deviant interaction with the neuronal receptor, Plexin B^28^.

Here, we extend our neuromuscular analyses in CMT2D, through a comprehensive assessment of developmental and degenerative processes at the mouse NMJ across several anatomically and functionally distinct wholemount muscles. By correlating varied neuropathic NMJ phenotypes with extensive morphological data on developing and mature wild-type synapses^29^, we have begun to identify key features underlying the selective vulnerability of neuromuscular connections in *GARS1* neuropathy.

## Materials and Methods

### Animals

Mouse work was carried out under license from the UK Home Office in accordance with the Animals (Scientific Procedures) Act 1986 and approved by the UCL Queen Square Institute of Neurology Ethical Review Committee. *Gars*^*C201R/+*^ mice (RRID: MGI 3849420) were maintained as heterozygote breeding pairs on a C57BL/6J background and genotyped as previously described^24^. Animals sacrificed for one month and three month timepoints were 27-35 and 84-95 days old, respectively, and both sexes were used unless otherwise stated.

### Grip strength testing

Grip strength was assessed in forelimbs as previously described^14^.

### Muscle dissection and immunohistochemistry

Muscles were dissected and stained as wholemount preparations as outlined formerly^30,31^. The following antibodies were used to co-stain pre-synaptic motor nerve terminals and axons: 1/25 mouse pan anti-synaptic vesicle 2 (SV2, Developmental Studies Hybridoma Bank [DSHB], Iowa City, IA, supernatant) and 1/250 mouse anti-neurofilament (2H3, DSHB, supernatant). 1/1,000 Alexa Fluor 555 α-BTX (Thermo Fisher Scientific, Waltham, MA, B35451, RRID:AB_2617152) was used to identify post-synaptic AChRs.

### NMJ imaging and analysis

NMJs were imaged using an inverted LSM780 laser scanning microscope (Zeiss, Oberkochen, Germany). Denervation, polyinnervation, and perforation analyses were performed as detailed elsewhere^25,30^.

### Statistical analysis

Data were assumed to be normally distributed unless evidence to the contrary could be provided by the D’Agostino and Pearson omnibus normality test, while equal variance between groups was assumed. GraphPad Prism 8 (version 8.4.0, La Jolla, CA) was used for all statistical tests, which were all two-sided. Datasets were statistically compared using either a one- or two-way analysis of variance (ANOVA) followed by either Sidak’s or Tukey’s multiple comparisons test. Correlation was assessed using Pearson’s product moment correlation. Bonferroni correction was applied to all correlation analyses in which multiple tests were performed in order to maintain an α of 0.05. Sample sizes, which were pre-determined using previous experience^25^ and power calculations, are reported in all figure legends and represent biological replicates (*i.e.* individual animals). Once stained and imaged, no muscle samples were excluded from analyses. As no treatment groups were involved, randomisation was not performed. All reported error bars depict standard error of the mean (SEM). All analyses were performed blinded to genotype.

## Results

### NMJ denervation underlies greater hindlimb weakness in CMT2D mice

Mutant *Gars* mice display a developmental perturbation of sensory neuron fate in hindlimb-innervating lumbar dorsal root ganglia (DRG)^14^, which is not present in forelimb-innervating cervical DRG^32^, suggestive of length-dependent phenotypes in CMT2D mice. To determine whether the neuromuscular system displays a similar pathological pattern, we performed grip strength testing of mild *Gars*^*C201R/+*^ mice^24^. Previous data indicate that muscle function is impaired at both one and three months in male and female *Gars*^*C201R/+*^ mice when all limbs are simultaneously tested^14^. Assessing just forelimb muscle function of female wild-type and *Gars*^*C201R/+*^ mice at the same timepoints, we observed a similar early and persistent deficit in CMT2D strength (**Supplementary Fig. S1**). Hindlimbs alone cannot be tested because of the manner in which mice grasp the grip strength meter used. Nevertheless, to compare grip strength from all limbs^14^ with forelimb data generated here, we normalised to wild-type at each timepoint in each limb category. The relative forelimb strength was significantly greater at both ages compared to all limbs (72% vs. 57% and 69% vs. 59%, **Fig. 1A**), suggesting that CMT2D mice may display greater weakness in hindlimbs.

**Figure 1.**
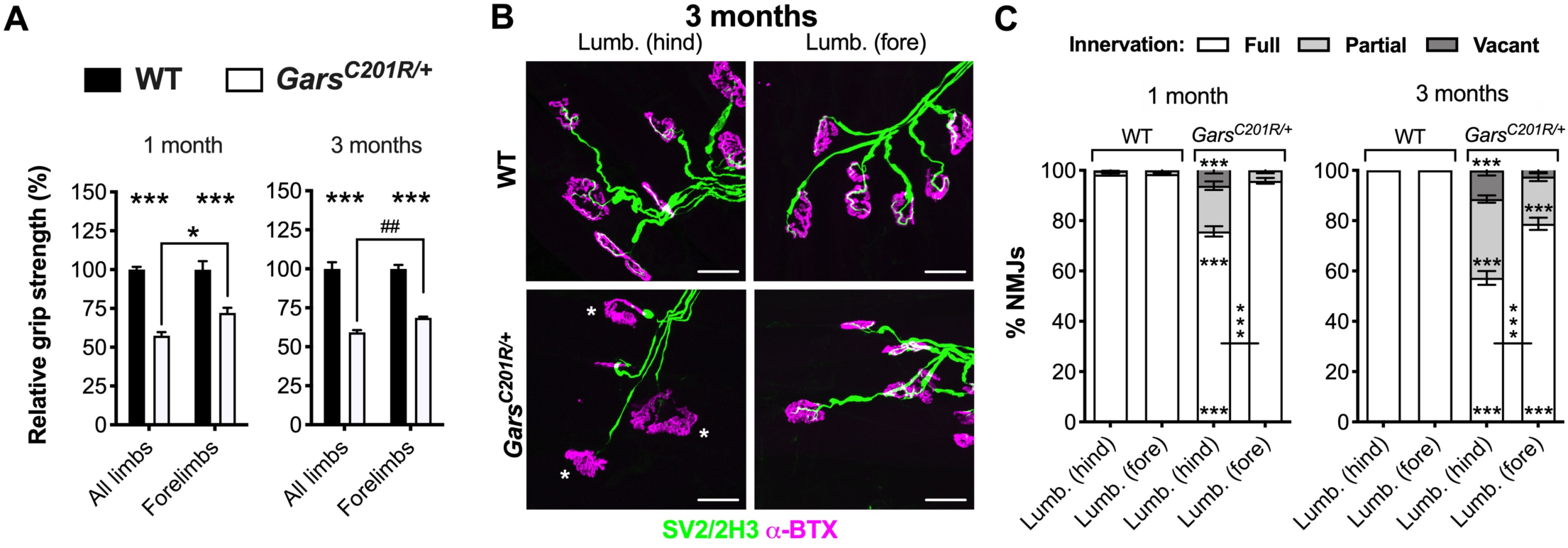
NMJ denervation underlies the greater weakness in CMT2D mouse hindlimbs. (**A**) Grip strength relative to wild-type [WT] is significantly impaired in all four limbs and just the forelimbs of *Gars*^*C201R/+*^ mice at one (left, genotype *P* < 0.001, limbs *P* = 0.049, interaction *P* = 0.049; two-way ANOVA) and three months (right, genotype *P* < 0.001, limbs *P* = 0.125, interaction *P* = 0.125; two-way ANOVA). The weakness is more pronounced when all limbs are tested than when just the forelimbs are assessed. *n* = 4-8. ## *P* < 0.01; unpaired *t*-test. (**B**) Representative collapsed z-stack confocal images of NMJs in hindpaw [Lumb. (hind)] and forepaw [Lumb. (fore)] lumbrical muscles dissected from three month old wild-type and *Gars*^*C201R/+*^ mice. Lower motor neurons are visualised using a combination of SV2/2H3 (green) and post-synaptic AChRs with α-BTX (magenta). Asterisks identify partially denervated synapses. Scale bars = 25 µm. (**C**) The hindpaw, but not forepaw, lumbrical muscles show significant denervation at one month. By three months, loss of innervation is worse in both muscles, with significant denervation now also observed in forepaw lumbricals. *P* < 0.001 for all one-way ANOVAs comparing the percentage of fully innervated, partially denervated and vacant NMJs in both muscles at both timepoints (six tests, three per age). *N.b.*, forepaw and hindpaw lumbricals were dissected from the same mice. *n* = 6. Means ± SEM are plotted in all graphs. *** *P* < 0.001, * *P* < 0.05; Sidak’s multiple comparisons test. See also **Supplementary Fig. S1**.

Lumbrical muscles involved in hindpaw clasping and grip strength, show a progressive loss of lower motor neuron connectivity in mutant *Gars* mice, correlating with overall model severity^25^. To compare NMJ innervation between a hindlimb and a forelimb muscle of similar function and relative limb location, we dissected both hindpaw and forepaw lumbrical muscles and stained them with antibodies against SV2/2H3 to visualise motor neurons and α-bungarotoxin (α-BTX) to identify post-synaptic acetylcholine receptors (AChRs) (**Fig. 1B**). At one month, hindlimb lumbricals displayed significantly more partially and fully (*i.e.* vacant) denervated NMJs than wild-type mice, becoming worse by three months (**Fig. 1C**). In contrast, forelimb lumbricals displayed no significant loss of innervation at one month, with comparatively mild denervation manifesting by three months. When statistically compared, loss of NMJ integrity was consistently significantly greater in hindpaw lumbricals, indicating that neuromuscular denervation contributes to the greater hindlimb weakness of mutant *Gars* mice.

### Impaired NMJ maturation precedes denervation

Prior to denervation, hindpaw lumbricals in CMT2D mice display impairments in key maturation events, such as synapse elimination and post-synaptic plaque-to-pretzel migration^25^; however, these developmental delays were not seen in a postural, anterior abdominal muscle called the transversus abdominis (TVA)^33^, which also lacked denervation at three months in mutant *Gars* mice^25^. To test whether this link between perturbed NMJ maturation and degeneration is consistent across CMT2D muscles, we assessed synaptic development in hindpaw and forepaw lumbricals (**Fig. 2**). At birth, post-synaptic endplates are contacted by several different motor neurons (see **Fig. 2A** for an example) and, through a process called synapse elimination, go from being polyinnervated to monoinnervated by two weeks in mice^34,35^. In tandem, AChRs migrate and become concentrated in close apposition to motor nerve terminals, resulting in their conversion from simple, circular plaques at birth to complex pretzel-like structures with increasing numbers of perforations^36^. Delayed synapse elimination was replicated in *Gars*^*C201R/+*^ hindpaw lumbrical NMJs, which showed significant polyinnervation at one month and three months (**Fig. 2B**). Contrastingly, forepaw lumbricals showed no such defect. To assess post-synaptic development, perforations per endplate were counted, with higher numbers indicative of advanced maturation. Once again, hindpaw lumbrical NMJs showed impaired AChR development, which was also present in forepaw lumbricals, albeit not to the same extent (**Fig. 2C**). These data indicate that defective pre-synaptic maturation is associated with subsequent neuromuscular degeneration, whereas the post-synaptic disruption may represent a systemic effect, as it is also seen in the non-denervated TVA^25^.

**Figure 2.**
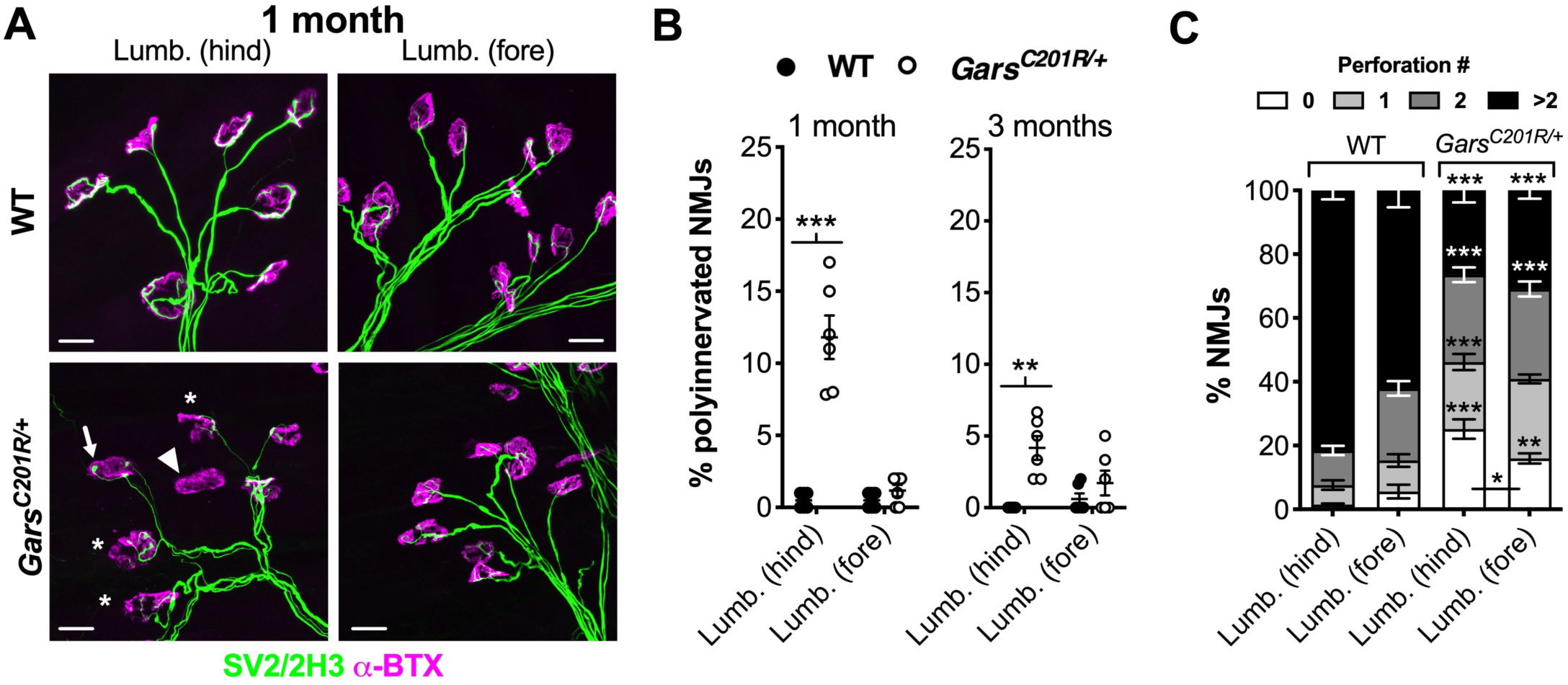
Impaired CMT2D NMJ maturation is linked to denervation. (**A**) Representative collapsed z-stack confocal images of NMJs in hindpaw [Lumb. (hind)] and forepaw [Lumb. (fore)] lumbrical muscles dissected from one month old *Gars*^*C201R/+*^ mice. Lower motor neurons are visualised using a combination of SV2/2H3 (green) and post-synaptic AChRs with α-BTX (magenta). Asterisks identify partially denervated synapses, the arrowhead a vacant synapse, and the arrow a polyinnervated NMJ. Scale bars = 25 µm. (**B**) Hindpaw, but not forepaw, lumbrical muscles of *Gars*^*C201R/+*^ mice display significantly more polyinnervated NMJs at one month compared to wild-type (left, genotype *P* < 0.001, muscle *P* < 0.001, interaction *P* < 0.001; two-way repeated measures ANOVA). This pattern is also present at three months (right, genotype *P* = 0.001, muscle *P* = 0.191, interaction *P* = 0.029; two-way repeated measures ANOVA), although there are fewer polyinnervated NMJs in the hindpaw lumbricals at this stage, indicative of a delay rather than cessation of synapse elimination. (**C**) Both hindpaw and forepaw lumbrical NMJs show post-synaptic maturation deficiency as assessed by counting the perforations per endplate. This defect was significantly worse in hindpaw lumbricals. *P* < 0.001 for all one-way ANOVAs comparing the percentage of NMJs with none, one, two, and more than two perforations in both muscles (*i.e.* four tests). *N.b.*, hindpaw and forepaw muscles are the same as those analysed in **Fig. 1**. *n* = 6. Means ± SEM are plotted in all graphs. *** *P* < 0.001, ** *P* < 0.01, * *P* < 0.05; Sidak’s multiple comparisons test.

### Hindlimb FDB muscles also display delayed NMJ development and degeneration

To extend our analyses into additional hindlimb and forelimb muscles, we stained NMJs of hindpaw flexor digitorum brevis (FDB) and forelimb epitrochleoanconeus (ETA) muscles dissected from three month old mice (**Fig. 3A**). FDB muscles aid hindpaw opening and the ETA contributes to forearm supination^37,38^, and both can be wholemount stained to assess innervation^29^. The FDB displayed defective synapse elimination associated with severe denervation, whereas the ETA was unaffected (**Fig. 3B-C**). These data uphold that CMT2D hindlimbs display greater pathology, and that impaired NMJ development consistently accompanies degeneration.

**Figure 3.**
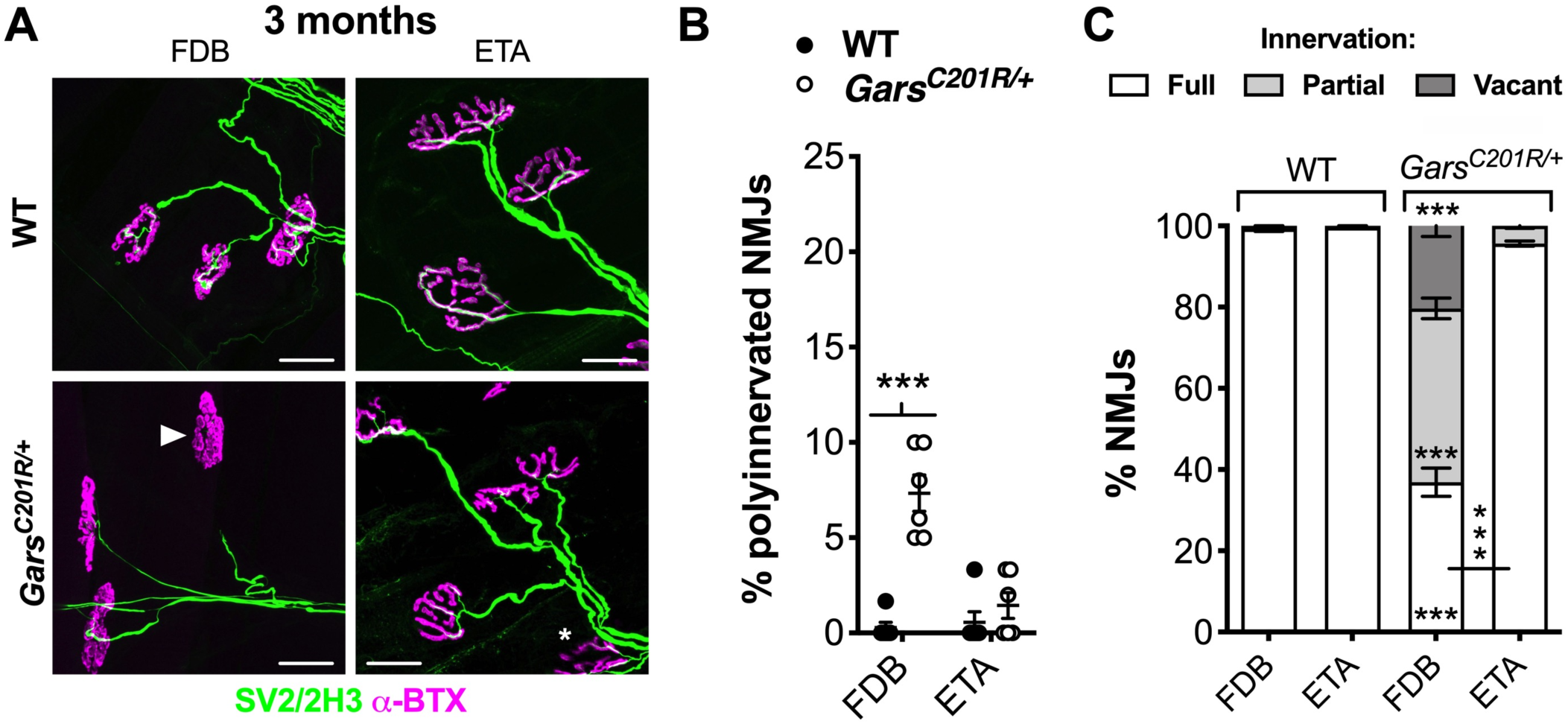
Hindlimb muscles behave similarly and show marked differences with forelimb muscles. (**A**) Representative collapsed z-stack confocal images of NMJs in FDB and ETA muscles dissected from three month old wild-type and *Gars*^*C201R/+*^ mice. Lower motor neurons are visualised using a combination of SV2/2H3 (green) and post-synaptic AChRs with α-BTX (magenta). The asterisk identifies a partially denervated NMJ and the arrowhead a fully denervated synapse. Scale bars = 25 µm. (**B**) The FDB, but not ETA, muscle of *Gars*^*C201R/+*^ mice displays significantly more polyinnervated NMJs at three months (genotype *P* < 0.001, muscle P = 0.005, interaction *P* < 0.001; two-way repeated measures ANOVA). (**C**) At three months, denervation in the FDB muscle is severe, whereas the ETA shows little degeneration. *P* < 0.001 for all one-way ANOVAs comparing the percentage of fully innervated, partially denervated and vacant NMJs in both muscles (*i.e.* three tests). *n* = 6. Means ± SEM are plotted in all graphs. *** *P* < 0.001; Sidak’s multiple comparisons test.

### CMT2D polyinnervation correlates with degeneration and is independent of synapse elimination

Including previously generated data from the TVA^25^, we have now assessed NMJ polyinnervation and denervation at three months in five wholemount muscles (**Fig. 4A**). We detected a broad spectrum of denervation (partial and full combined) in CMT2D mice, ranging from 2% in the TVA to 63% in the FDB (**Fig. 4B, Supplementary Table S1**). A similar phenotypic continuum was identified for disturbed synapse elimination, with 0.7% of TVA NMJs remaining polyinnervated and up to 7.3% of FDB NMJs (**Fig. 4C, Supplementary Table S1**). We tested the relationship between these two neuropathic phenotypes and found a highly significant positive correlation between percentage of vacant and polyinnervated synapses occurring at three months (**Fig. 4D**). A similar correlation was present when vacant and partially denervated synapses were combined (% denervation, **Fig. 4E**). To determine whether perturbed synapse elimination is simply a delay of the physiological process occurring in wild-type animals, we correlated the CMT2D motor input counts with previously generated^29^ P7 wild-type polyinnervation data from the five wholemount muscles (**Fig. 4F**). No correlation was identified, suggesting that impaired synapse elimination in mutant *Gars* mice may be caused by selective, muscle-specific perturbation of pathways relevant to removal of supernumerary motor axons, as opposed to a systemic slow-down in the process.

**Figure 4.**
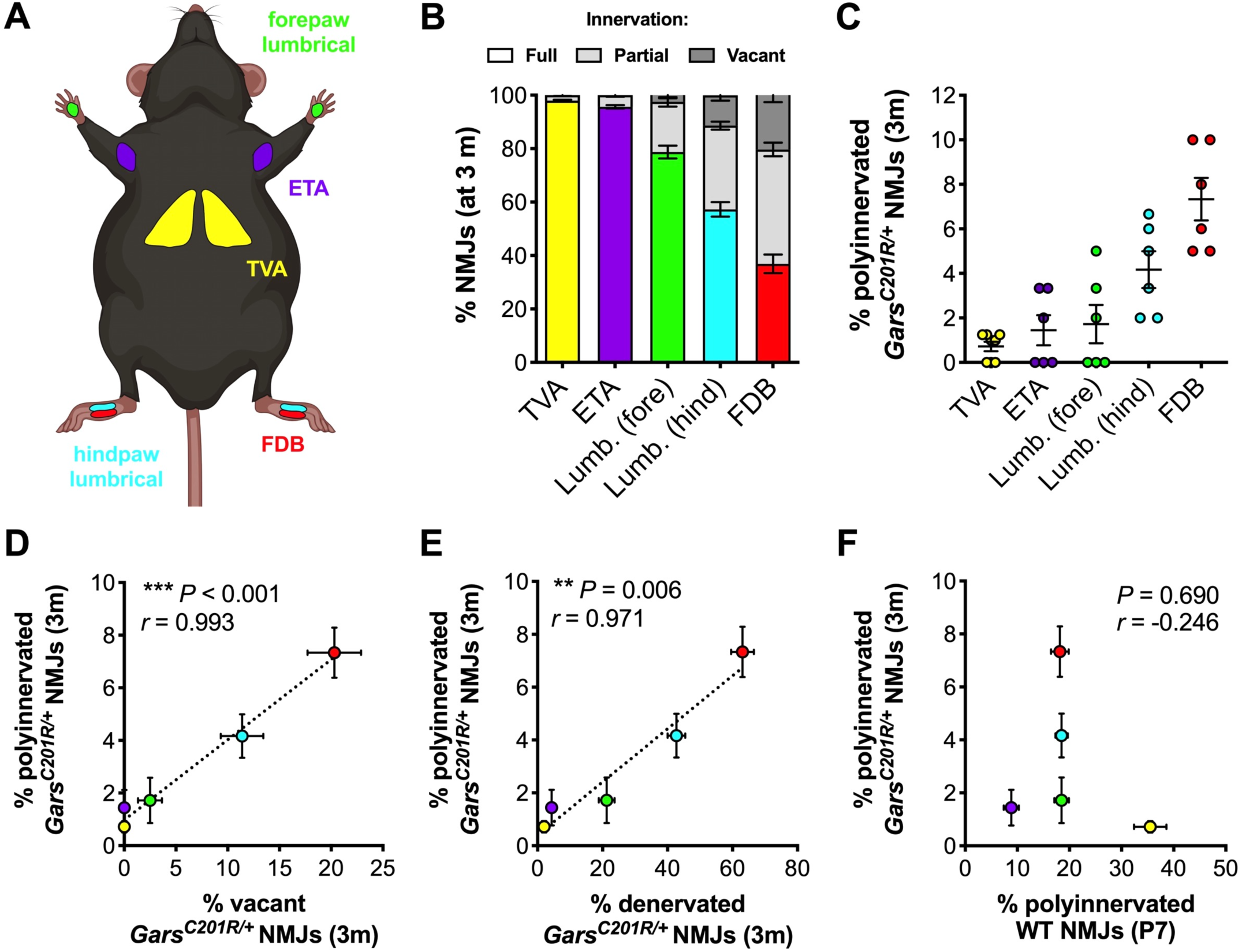
CMT2D NMJ polyinnervation correlates with denervation and is not simply a delay of wild-type synapse elimination. (**A**) Five thin and flat muscles were analysed in this study: transversus abdominis (TVA, yellow), epitrochleoanconeus (ETA, purple), forelimb lumbricals (green, Lumb. [fore]), hindlimb lumbricals (cyan, Lumb. [hind]) and flexor digitorum brevis (FDB, red). The schematic was created with BioRender (https://biorender.com). (**B**) The five muscles display a spectrum of vulnerability to denervation in *Gars*^*C201R/+*^ mice at three months; the TVA remains unaffected, while the FDB muscles show stark degeneration. (**C**) Defective synapse elimination at three months is more pronounced in muscles with greater denervation. (**D, E**) There is a significant correlation between the percentage of polyinnervation observed in each muscle and the percentage of vacant (D, *** *P* < 0.001, *r* = 0.993) or denervated (E, ** *P* = 0.006, *r* = 0.971) NMJs at three months in *Gars*^*C201R/+*^ mice. (**F**) There is no correlation between the percentage of wild-type NMJ polyinnervation at P7 and the polyinnervation percentage at three months in *Gars*^*C201R/+*^ mice (*P* = 0.690, *r* = - 0.246), suggesting that the mutant phenotype is not due to a delay of wild-type synapse elimination. See **Supplementary Table S1** for pairwise statistical testing between muscles in panels B and C. Data in panels E-F were analysed by Pearson’s product moment correlation. The colour-coding of muscles in panels E-F is maintained from panels A-C. TVA data^25^ and wild-type P7 polyinnervation data^29^ were generated for previous studies. *n* = 6-8. Means ± SEM are plotted in all graphs.

### Fibre type does not associate with CMT2D denervation

Motor neurons innervating hindpaw muscles (FDB and lumbricals) are more impacted in CMT2D mice than those targeting forepaws (lumbricals) followed by more proximal muscles (ETA and TVA) (**Fig. 4**). This indicates a general distal-to-proximal vulnerability axis consistent with length-dependent axon degeneration. However, adjacent muscles innervated by neurons of similar length show significant distinctions in denervation (**Supplementary Table S1**), suggesting that additional factors contribute to NMJ vulnerability. Fast-fatiguable motor neurons innervating fast twitch muscle fibres are more susceptible to degeneration in amyotrophic lateral sclerosis (ALS) than motor nerves contacting slow twitch fibres^39^. To assess whether this is also observed in CMT2D, we correlated previously published fast twitch fibre type percentages^29,37,38,40–42^ with mutant *Gars* mouse denervation (**Supplementary Fig. S2**). No relationship was observed indicating that muscle fibre type is unlikely to be driving inter-muscle disparities in neuropathy.

### CMT2D denervation correlates with demand for post-natal NMJ growth

We then evaluated NMJ structure as a possible determinant in neuropathology, since NMJs with smaller synaptic volumes have less neuronal input to lose to become fully denervated. Using a robust and standardised semi-automated workflow called NMJ-morph^43^, we have previously generated detailed data on morphology of developing (P7) and mature (P31-32) wild-type NMJs in the same five muscles assessed here^29^. These datasets were used to show that developing synapses show greater inter-muscle variability and that post-natal NMJ growth occurs at different rates across muscles^29^ – two phenotypes that may provide a substrate for differential NMJ degeneration in CMT2D. We therefore first correlated all 41 morphological variables from P7 and P31-32 wild-type NMJs with the percentage of vacant synapses in three month mutant *Gars* mice (**Supplementary Table S2**); however, no significant relationships were identified. We then tested for correlations between morphology and the combined percentages of partially and fully denervated NMJs (**Supplementary Table S3**). Of the 41 variables, only AChR perimeter of developing NMJs showed a significant interaction, indicating that a smaller AChR perimeter is associated with greater levels of denervation (**Supplementary Fig. S3**). Given that many of the assessed NMJ variables are interrelated (e.g. AChR perimeter and area), the finding that only one out of 82 tests showed a significant correlation suggests that synaptic architecture has relatively little impact on the extent of neuropathy.

Finally, we assessed the role of post-natal NMJ growth and development by correlating the percentage change in each morphological variable with the extent of mutant *Gars* degeneration. Several different pre-synaptic (terminal area, number of branches and branch length), post-synaptic (AChR area) and overlapping (synaptic contact area) variables significantly correlated with the percentage of vacant CMT2D NMJs (**Fig. 5A, Supplementary Table S4**). Two further pre-synaptic variables (branch point number and complexity) significantly associated with the percentage of denervated (partial and full) mutant NMJs (**Fig. 5B, Supplementary Table S5**). All seven correlations were positive, indicating that NMJs undergoing greater post-natal change in morphology are more prone to degeneration in CMT2D mice.

**Figure 5.**
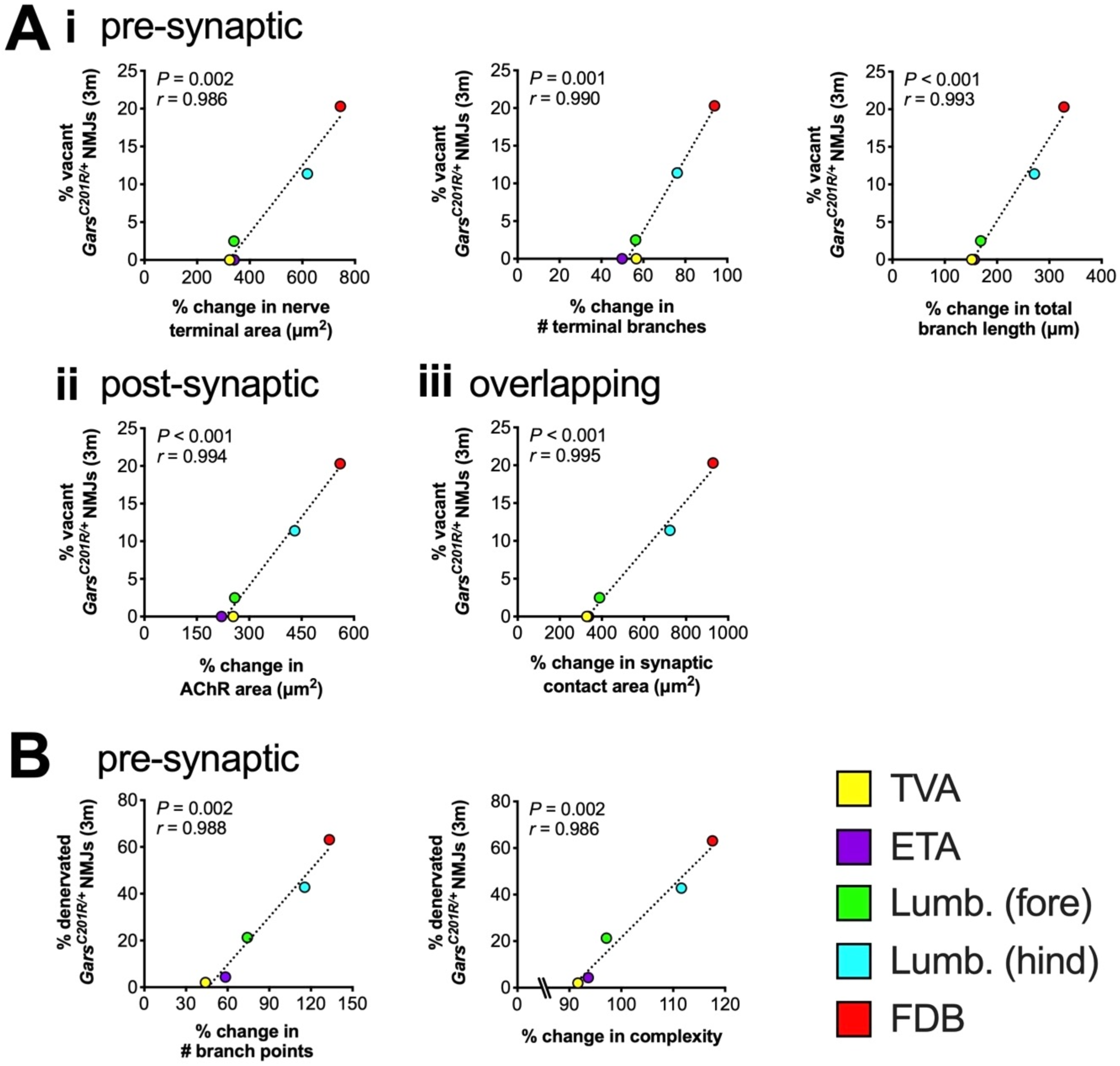
NMJs that undergo greater post-natal morphological change are more vulnerable to degeneration in CMT2D mice. The percentages of vacant (**A**) and denervated (**B**) NMJs in three month *Gars*^*C201R/+*^ mice were correlated with post-natal NMJ growth in 20 different pre-synaptic (i), post-synaptic (ii) and overlapping (iii) morphological variables. Wild-type P7 to P31-32 morphological data^29^ and TVA vacancy data^25^ were previously published. Correlation was assessed by calculating Pearson’s product moment correlation coefficient (*r*), the results of which are presented in **Supplementary Table S4** and **Supplementary Table S5** along with associated *P* values. Only significant correlations are presented, *i.e.* Bonferroni-corrected *P* < 0.00256. *n* = 6 (all muscles except TVA) and 8 (TVA). Lumb. (fore), forepaw lumbricals; Lumb. (hind), hindpaw lumbricals.

## Discussion

CMT patients usually present with weakness and atrophy in the feet and then hands, indicating that peripheral neurons with longer axons are generally more impacted by the disease. However, mutations in a few genes, including *GARS1*, can cause an upper limb predominant phenotype^7,8,44^. This suggests that while axon length is important, additional neuronal characteristics contribute to pathogenesis and the spectrum of motor neuron involvement. Identification of such factors may elucidate pathomechanisms and provide potential targetable pathways. While we did not observe upper limb weakness in mutant *Gars* mice (possible reasons for which are discussed elsewhere in relation to sensory phenotypes^32^), we do observe significant distinctions in NMJ pathology between adjacent muscles innervated by motor neurons of similar length. This suggests that, like in human patients, CMT2D mouse motor neurons possess features that modify their predisposition to neuropathology.

We first assessed the impact of muscle fibre type on neuropathy, because large, fast-fatiguable motor neurons targeting fast twitch fibres are more vulnerable in ALS^39^, a motor neuron disease with parallels to CMT. There was no correlation between fibre type and CMT2D degeneration, suggesting that in contrast to ALS, muscle fibre types have little impact on the extent of motor neuropathy. Since pathology in mutant *Gars* mice and patients is restricted to distal segments of peripheral nerves, we then probed NMJ morphology. Again, we found little evidence to indicate that distinctions in pre- or post-synaptic architecture cause the spectrum of synapse loss. However, we did find that CMT2D denervation correlated closely with wild-type NMJ change and development. This suggests that motor terminals under the greatest pressures to grow and mature during early post-natal life, perhaps due to different functions of their associated muscles^29^, are more likely to degenerate. Possibly contributing to this, severe CMT2D mice possess fewer mitochondria in pre-synaptic motor terminals in the proximal levator auris longus muscle^26^. Accordingly, motor neurons with higher bioenergetic capacity and thus energy supply have been shown to be more resistant to disease in a mouse model for spinal muscular atrophy^45^, while computational modelling indicates energetic demand also contributes to selective vulnerability in ALS^46^.

While developmental demand may contribute to early synaptic disruption, *GARS1*-associated neuropathy is progressive and denervation continues in CMT2D mice up to three months and beyond. Additional mechanisms are therefore likely driving the life-long neurodegeneration caused by mutant GlyRS.

Synapse elimination is mediated by reciprocal nerve-muscle signalling and electrical activity to refine nervous system architecture^35,47^. The strong correlation across CMT2D muscles between polyinnervated and denervated synapses, and the lack of association with wild-type polyinnervation, indicate that synapse elimination is linked to pathology and not caused by systemic interruption.

Synapse elimination delay is unlikely to be driving denervation for two reasons; firstly, having multiple motor neurons innervating an endplate is probably beneficial from an innervation standpoint as several motor nerve terminals must degenerate for full denervation to occur. Corroborating this, the percentage of denervation (partial and full) across all FDB NMJs is 63 ± 4% at three months, whereas at polyinnervated synapses, only 6 ± 4% of NMJs show signs of degeneration (data not shown). Secondly, the mechanisms mediating removal of supernumerary motor inputs share many features with neurodegenerative pathways^48^, and hence a delay in elimination would likely also postpone neuropathy. Alternatively, the phenotype could represent a compensatory response to degeneration, similar in essence to neuronal sprouting aided by terminal Schwann cells^49^. However, if this were the case, it would probably be observed throughout the lifespan of CMT2D mice and in different neuromuscular disease models, but this does not appear to occur. We therefore suggest that the polyinnervation phenotype is an epiphenomenon resulting from aberrant interaction of mutant GlyRS with (a) key trans-synaptic protein(s). There is precedent for this in a CMT2D *Drosophila* model; muscle-secreted mutant GlyRS aberrantly accumulates at NMJs prior to degeneration^16^ and coincides with neurodevelopmental wiring defects^28^. This phenocopies an ectopic motor neuron branching caused by loss-of-function mutations in *plexA* and *plexB*, which encode neuronal transmembrane proteins that bind secreted semaphorins to facilitate axonal guidance, retrograde signalling and synaptic plasticity^50,51^. This observation led to the finding that plexin B serves as an erroneous binding partner for mutant GlyRS at the NMJ; however, while mutant GlyRS interferes with the fidelity of axon guidance pathways causing wiring defects, this was de-coupled from the NMJ pathologies, *i.e.* was an epiphenomenon^28^.

We have previously shown that mutant GlyRS aberrantly interacts with neurotrophin receptors (TrkA-C)^14^, of which TrkB is the main one present at the mammalian NMJ^52,53^. TrkB and its ligand, brain-derived neurotrophic factor (BDNF), play an important role in synapse elimination at the NMJ^53–55^. This may, in part, be driven by BDNF-mediated clustering in motor nerve terminals of voltage-gated calcium channels, which facilitate calcium influx, synaptic neurotransmission, and NMJ maturation^56–59^. Mutant GlyRS mis-interacting with TrkB at the NMJ could delay synapse elimination through impairing calcium channel insertion into pre-synaptic motor terminals, thereby hindering neurotransmission and neuronal activity. Consistent with this idea, it has been shown that CMT2D mice display pre-synaptic transmission defects, including reduced frequency of spontaneous neurotransmitter release, which is calcium transient-dependent^26^. A primary mechanism causing the weakened synaptic transmission was ultimately not identified, instead it was suggested that several different release processes function sub-optimally^26^. It is therefore conceivable that accumulation of mutant GlyRS at NMJs through association with TrkB, akin to the synaptic build-up of pathological protein observed in the fly model^16,28^, may be contributing to the impaired synaptic function prior to NMJ degeneration.

Upon BDNF binding, TrkB is internalised into neuron terminals. Activated receptor complexes can then precipitate local signalling events such as those just discussed or become sorted into signalling endosomes for long-range retrograde axonal transport critical to pro-survival gene transcription^60,61^. The erroneous association of mutant GlyRS with TrkB at motor nerve terminals may thus diminish retrograde neurotrophic signalling, contributing to neuropathy. Correspondingly, signalling endosome transport is impaired in primary sensory neurons cultured from severe mutant *Gars* mice^62^, while mutations in essential signalling endosome retrograde effectors, Rab7 (Ref. 63) and dynactin, cause peripheral neuropathy^64,65^. Additionally, activation of IGF1R, a neurotrophic factor receptor found at the NMJ, is involved in regulating signalling endosome dynamics in motor neurons^66^, which may be a common function of similar receptors like TrkB^67^. Furthermore, high levels of phosphorylated IGF1R at NMJs may confer resistance to motor degeneration in ALS mice^68^, suggesting that levels of neurotrophic factors and/or their receptors (e.g. BDNF/TrkB) could indeed play a role in selective vulnerability to motor terminal loss in peripheral neuropathy. Alternatively, mutant GlyRS sequestration of TrkB could divert BDNF to bind to the pan-neurotrophin death receptor, p75NTR, resulting in over-activation of degenerative signals^69^. Consistent with these views and the importance of TrkB signalling to peripheral synapse integrity, reduced TrkB activity is known to impair NMJ neurotransmission, maturation and innervation, without impacting motor neuron number^52,70^, phenotypes very similar to those found in CMT2D mice. It is therefore conceivable that different levels of TrkB and/or BDNF at NMJs across muscles contribute to both the differential denervation and subverted synaptic maturation.

In summary, by assessing developmental and degenerative NMJ phenotypes across five different wholemount muscles from CMT2D mice, we have identified a spectrum of vulnerability to neurodegeneration. Through correlation analyses with muscle fibre type, NMJ architecture, and post-natal synaptic growth, we identified that the magnitude of developmental demand likely contributes to neuromuscular synapse loss in mutant *Gars* mice. We believe that these results point the way forward for an improved understanding of the molecular mechanisms driving differences in synaptic vulnerability to neuropathy.

## Acknowledgements

The authors would like to thank Stuart J. Grice (University of Oxford) and David Villarroel-Campos (University College London) for critical comments on the manuscript and Anna-Leigh Brown (University College London) for fruitful discussions. This work was supported by the Medical Research Council Career Development Award (MR/S006990/1) [JNS], the Wellcome Trust Sir Henry Wellcome Postdoctoral Fellowship (103191/Z/13/Z) [JNS], the Wellcome Trust Senior Investigator Award (107116/Z/15/Z) [GS], the European Union’s Horizon 2020 Research and Innovation programme under grant agreement 739572 [GS], and the UK Dementia Research Institute Foundation award (UKDRI-1005) [GS].

## Author Contributions

JNS conceived the experiments; JNS, AMM performed the research; JNS, AMM analysed the data; GS provided expertise and discussion; JNS wrote the manuscript with input from all authors. All authors approved submission of this work.

## Supplementary Figures

**Supplementary Figure S1.**
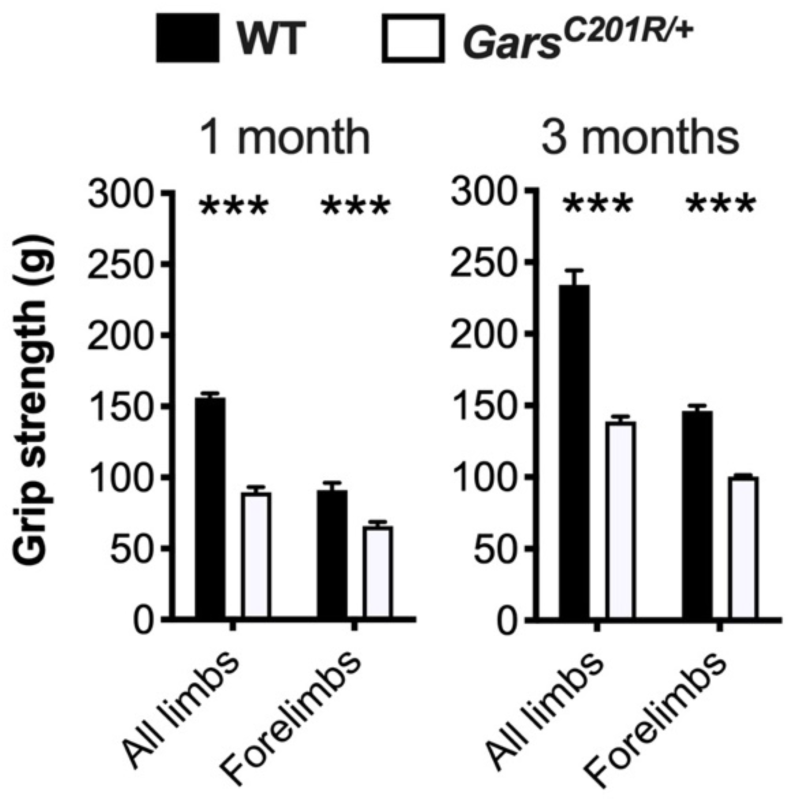
Raw grip strength data. Grip strength of all limbs and just the forelimbs are both significantly impaired in *Gars*^*C201R/+*^ mice at one (left, genotype *P* < 0.001, limbs *P* < 0.001, interaction *P* < 0.001; two-way ANOVA) and three months (right, genotype *P* < 0.001, limbs *P* < 0.001, interaction *P* < 0.001; two-way ANOVA). *N.b.*, only females were analysed, “All limbs” data has been previously published^1^, and different mice were used for the two timepoints and for testing all limbs vs. forelimbs. *n* = 4-8. Means ± SEM are plotted. ^***^*P* < 0.001; Sidak’s multiple comparisons test. See also **Fig. 1**.

**Supplementary Figure S2.**
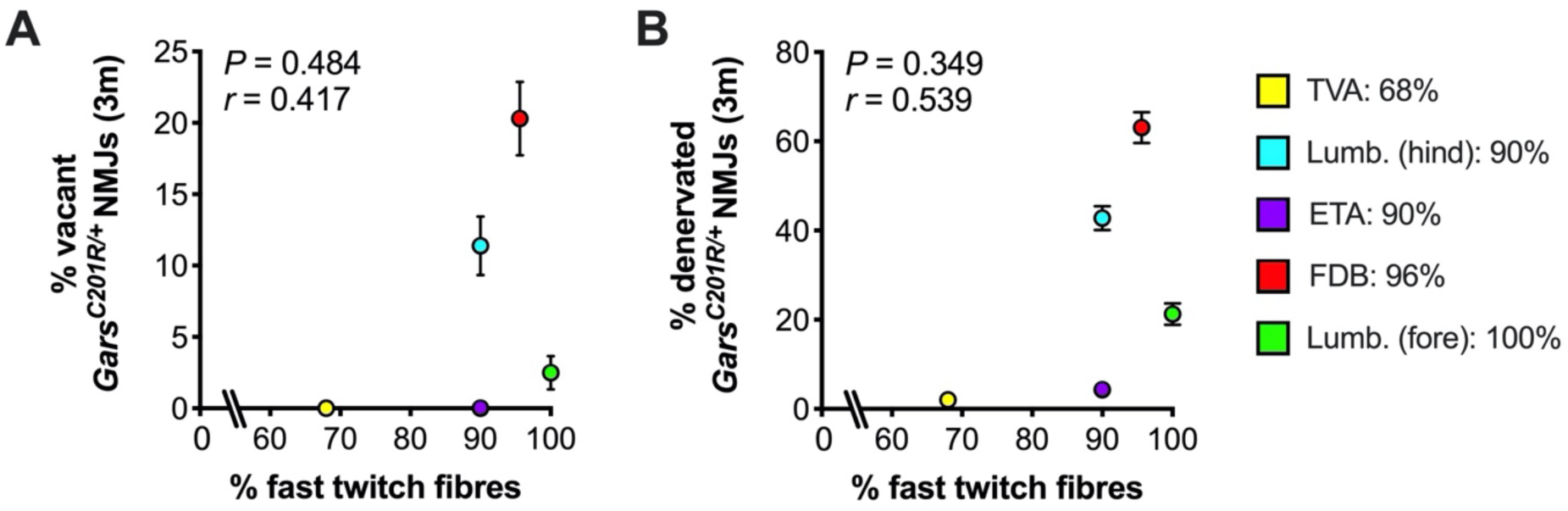
CMT2D denervation does not correlate with muscle fibre type. (**A, B**) There is no correlation between fast twitch fibres percentage found in each muscle (see legend for %) and percentage of vacant (A, *P* = 0.484, *r* = 0.417, Pearson’s product moment correlation) or denervated (B, *P* = 0.349, *r* = 0.539, Pearson’s product moment correlation) NMJs at three months in *Gars*^*C201R/+*^ mice. *n* = 6. Means ± SEM are plotted for % vacancy (A) and % denervation (B) in CMT2D mice. *Lumb. (fore)*, forepaw lumbricals; *Lumb. (hind)*, hindpaw lumbricals. See also **Fig. 4**.

**Supplementary Figure S3.**
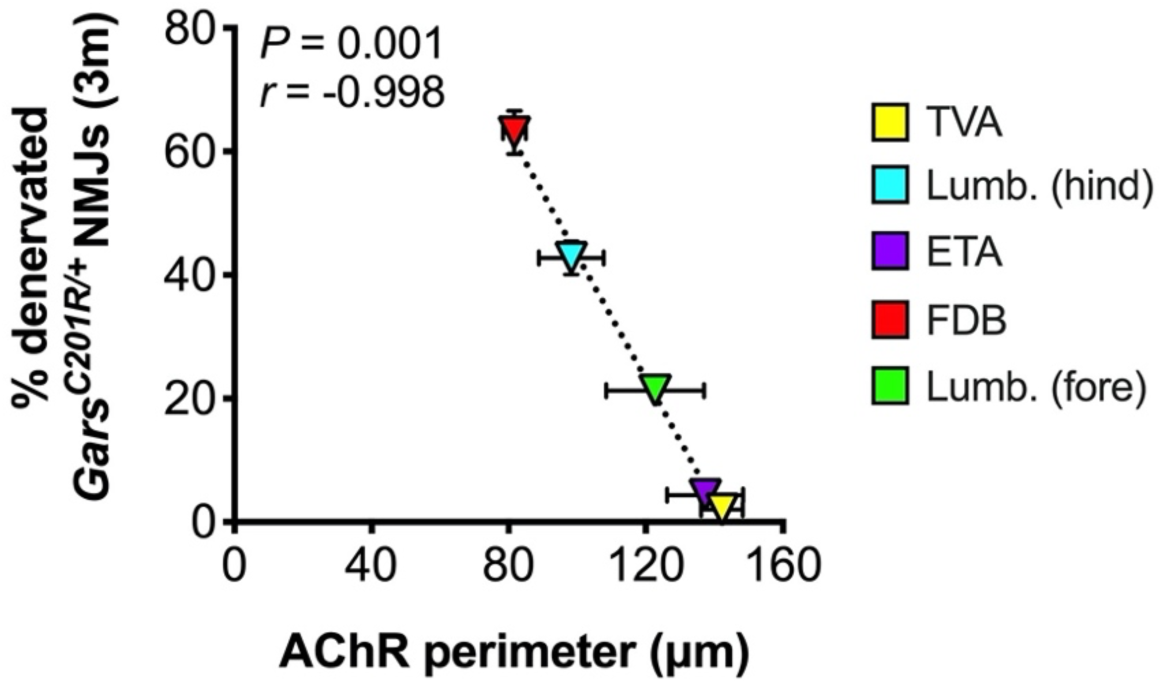
AChR perimeter of immature NMJs is the only morphological variable to correlate with CMT2D denervation. Correlation was assessed between the percentage of vacant or denervated NMJs in three month *Gars*^*C201R/+*^ mice and 21 different NMJ morphological variables from wild-type mice at both P7 and P31-32. Wild-type morphological data^2^ and TVA vacancy/denervation data^3^ have been published elsewhere. Correlation was assessed by calculating Pearson’s product moment correlation coefficient (*r*), the results of which are presented in **Supplementary Table S2** (% vacant) and **Supplementary Table S3** (% denervated) along with associated *P* values. Of 82 different tests, only AChR perimeter at P7 significantly correlated (Bonferroni-corrected *P* < 0.00244 for the 21 tests per timepoint) with the percentage denervation at the CMT2D NMJ (*P* = 0.001, *r* = −0.998). *n* = 6-8. Means ± SEM are plotted. *Lumb. (fore)*, forepaw lumbricals; *Lumb. (hind)*, hindpaw lumbricals.

## Supplementary Tables

**Supplementary Table S1.**
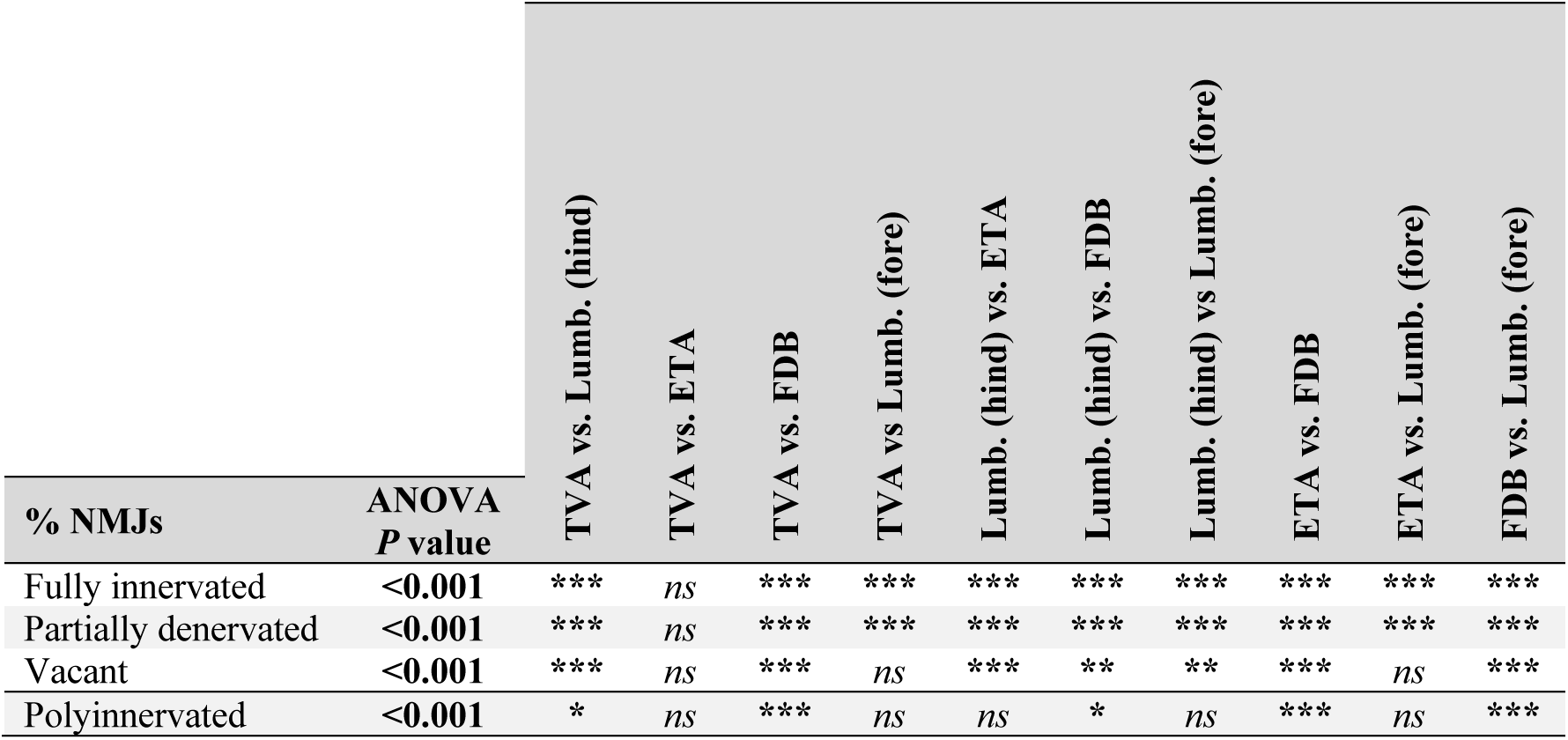
Pairwise significance testing between CMT2D mouse muscle denervation and polyinnervation. **\*\*\*** *P* < 0.001, ** *P* < 0.01, * *P* < 0.05, *ns* not significant; Tukey’s multiple comparisons test. *Lumb. (fore)*, forepaw lumbricals; *Lumb. (hind)*, hindpaw lumbricals. See also **Fig. 4**.

**Supplementary Table S2.** Statistical testing of correlation between the percentage of vacant *Gars*^*C201R/+*^ NMJs at three months and wild-type NMJ morphological variables at P7 and P31-32. Pre-synaptic variables are shaded green, post-synaptic variables shaded purple, and combined pre- and post-synaptic variables are unshaded. *n/a*, not applicable as wild-type morphological data not available at P7. See also **Supplementary Fig. S3**.

|  | P7 |  | P31-32 |  |
| --- | --- | --- | --- | --- |
| Variable | *r* | Pearson<br>*P* value | *r* | Pearson<br>*P* value |
| Polyinnervation (%) | -0.160 | 0.800 | -0.538 | 0.350 |
| Nerve terminal perimeter (μm) | -0.896 | 0.040 | -0.499 | 0.393 |
| Nerve terminal area (μm<sup>2</sup>) | -0.932 | 0.021 | -0.447 | 0.451 |
| # terminal branches | -0.769 | 0.128 | -0.490 | 0.404 |
| # branch points | -0.825 | 0.086 | -0.194 | 0.767 |
| Total branch length (μm) | -0.974 | 0.053 | -0.463 | 0.433 |
| Average branch length (μm) | -0.137 | 0.826 | 0.038 | 0.951 |
| Complexity | -0.883 | 0.047 | -0.294 | 0.632 |
| Axon diameter (μm) | -0.757 | 0.139 | -0.563 | 0.334 |
| AChR perimeter (μm) | -0.978 | 0.004 | -0.451 | 0.446 |
| AChR area (μm<sup>2</sup>) | -0.884 | 0.046 | -0.427 | 0.473 |
| Endplate diameter (μm) | -0.718 | 0.172 | -0.362 | 0.550 |
| Endplate perimeter (μm) | -0.840 | 0.075 | -0.447 | 0.451 |
| Endplate area (μm<sup>2</sup>) | -0.900 | 0.038 | -0.498 | 0.393 |
| # AChR clusters | -0.930 | 0.022 | -0.613 | 0.272 |
| AChR cluster area (μm<sup>2</sup>) | -0.809 | 0.097 | 0.846 | 0.071 |
| Compactness (%) | -0.915 | 0.029 | 0.730 | 0.162 |
| Fragmentation | -0.990 | 0.038 | -0.652 | 0.233 |
| Muscle fibre diameter (μm) | *n/a* | *n/a* | -0.441 | 0.457 |
| Synaptic contact area (μm<sup>2</sup>) | -0.920 | 0.027 | -0.450 | 0.447 |
| Overlap (%) | -0.889 | 0.044 | 0.216 | 0.728 |

**Supplementary Table S3.** Statistical testing of correlation between the percentage of denervated *Gars*^*C201R/+*^ NMJs at three months and wild-type NMJ morphological variables at P7 and P31-32. Pre-synaptic variables are shaded green, post-synaptic variables shaded purple, and combined pre- and post-synaptic variables are unshaded. ^***#***^*P* < 0.00244 Pearson’s product moment correlation, *i.e.* the Bonferroni correction-adjusted *P* value for an α of 0.05 when performing 21 associated tests. *n/a*, not applicable as wild-type morphological data not available at P7. See also **Supplementary Fig. S3**.

|  | P7 |  | P31-32 |  |
| --- | --- | --- | --- | --- |
| Variable | *r* | Pearson<br>*P* value | *r* | Pearson<br>*P* value |
| Polyinnervation (%) | -0.214 | 0.730 | -0.611 | 0.274 |
| Nerve terminal perimeter (μm) | -0.963 | 0.009 | -0.641 | 0.244 |
| Nerve terminal area (μm<sup>2</sup>) | -0.982 | 0.003 | -0.573 | 0.313 |
| # terminal branches | -0.873 | 0.053 | -0.641 | 0.244 |
| # branch points | -0.915 | 0.029 | -0.281 | 0.647 |
| Total branch length (μm) | -0.949 | 0.014 | -0.603 | 0.282 |
| Average branch length (μm) | 0.017 | 0.978 | 0.221 | 0.721 |
| Complexity | -0.956 | 0.011 | -0.447 | 0.450 |
| Axon diameter (μm) | -0.647 | 0.238 | -0.604 | 0.281 |
| AChR perimeter (μm) | -0.998 | **0.001<sup>#</sup>** | -0.601 | 0.283 |
| AChR area (μm<sup>2</sup>) | -0.956 | 0.011 | -0.571 | 0.315 |
| Endplate diameter (μm) | -0.814 | 0.093 | -0.491 | 0.401 |
| Endplate perimeter (μm) | -0.922 | 0.026 | -0.687 | 0.299 |
| Endplate area (μm<sup>2</sup>) | -0.964 | 0.008 | -0.639 | 0.246 |
| # AChR clusters | -0.864 | 0.059 | -0.745 | 0.149 |
| AChR cluster area (μm<sup>2</sup>) | -0.903 | 0.036 | 0.901 | 0.037 |
| Compactness (%) | -0.973 | 0.005 | 0.845 | 0.071 |
| Fragmentation | -0.844 | 0.072 | -0.778 | 0.121 |
| Muscle fibre diameter (μm) | *n/a* | *n/a* | -0.571 | 0.315 |
| Synaptic contact area (μm<sup>2</sup>) | -0.978 | 0.004 | -0.567 | 0.319 |
| Overlap (%) | -0.819 | 0.090 | 0.383 | 0.524 |

**Supplementary Table S4.** Statistical testing of correlation between the percentage of vacant *Gars*^*C201R/+*^ NMJs at three months and the percentage change in wild-type NMJ morphological variables from P7 to P31-32. Pre-synaptic variables are shaded green, post-synaptic variables shaded purple, and combined pre- and post-synaptic variables are unshaded. ^***#***^*P* < 0.00256 Pearson’s product moment correlation, *i.e.* the Bonferroni correction-adjusted *P* value for an α of 0.05 when performing 20 associated tests. See also **Fig. 5**.

| Variable | P7 to P31-32 |  |
| --- | --- | --- |
|  | *r* | Pearson *P* value |
| Polyinnervation (%) | -0.257 | 0.677 |
| Nerve terminal perimeter (μm) | 0.968 | 0.007 |
| Nerve terminal area (μm<sup>2</sup>) | 0.986 | **0.002<sup>#</sup>** |
| # terminal branches | 0.990 | **0.001<sup>#</sup>** |
| # branch points | 0.964 | 0.008 |
| Total branch length (μm) | 0.993 | **<0.001<sup>#</sup>** |
| Average branch length (μm) | 0.289 | 0.637 |
| Complexity | 0.982 | 0.003 |
| Axon diameter (μm) | -0.069 | 0.912 |
| AChR perimeter (μm) | 0.643 | 0.242 |
| AChR area (μm<sup>2</sup>) | 0.994 | **<0.001<sup>#</sup>** |
| Endplate diameter (μm) | 0.956 | 0.011 |
| Endplate perimeter (μm) | 0.943 | 0.016 |
| Endplate area (μm<sup>2</sup>) | 0.955 | 0.012 |
| # AChR clusters | -0.215 | 0.728 |
| AChR cluster area (μm<sup>2</sup>) | 0.928 | 0.023 |
| Compactness (%) | 0.882 | 0.048 |
| Fragmentation | -0.398 | 0.507 |
| Synaptic contact area (μm<sup>2</sup>) | 0.995 | **<0.001<sup>#</sup>** |
| Overlap (%) | 0.833 | 0.080 |

**Supplementary Table S5.** Statistical testing of correlation between the percentage of denervated *Gars*^*C201R/+*^ NMJs at three months and the percentage change in wild-type NMJ morphological variables from P7 to P31-32. Pre-synaptic variables are shaded green, post-synaptic variables shaded purple, and combined pre- and post-synaptic variables are unshaded. ^***#***^*P* < 0.00256 Pearson’s product moment correlation, *i.e.* the Bonferroni correction-adjusted *P* value for an α of 0.05 when performing 20 associated tests. See also **Fig. 5**.

| Variable | P7 to P31-32 |  |
| --- | --- | --- |
|  | *r* | Pearson *P* value |
| Polyinnervation (%) | -0.207 | 0.738 |
| Nerve terminal perimeter (μm) | 0.919 | 0.028 |
| Nerve terminal area (μm<sup>2</sup>) | 0.962 | 0.009 |
| # terminal branches | 0.962 | 0.009 |
| # branch points | 0.988 | **0.002<sup>#</sup>** |
| Total branch length (μm) | 0.977 | 0.004 |
| Average branch length (μm) | 0.460 | 0.435 |
| Complexity | 0.986 | **0.002<sup>#</sup>** |
| Axon diameter (μm) | -0.190 | 0.760 |
| AChR perimeter (μm) | 0.496 | 0.395 |
| AChR area (μm<sup>2</sup>) | 0.968 | 0.007 |
| Endplate diameter (μm) | 0.948 | 0.014 |
| Endplate perimeter (μm) | 0.896 | 0.040 |
| Endplate area (μm<sup>2</sup>) | 0.896 | 0.040 |
| # AChR clusters | -0.388 | 0.519 |
| AChR cluster area (μm<sup>2</sup>) | 0.980 | 0.003 |
| Compactness (%) | 0.954 | 0.012 |
| Fragmentation | -0.554 | 0.332 |
| Synaptic contact area (μm<sup>2</sup>) | 0.979 | 0.004 |
| Overlap (%) | 0.883 | 0.047 |

